# Network-based integration of cross-dataset proteomic profiles using fold-change directionality

**DOI:** 10.64898/2026.04.19.718092

**Authors:** Manaka Nishizaki, Norie Araki, Shin Kawano

## Abstract

**Motivation:** The rapid expansion of proteomic data has created new opportunities for large-scale integrative analyses. However, substantial variability across platforms, experimental designs, and processing pipelines limits direct quantitative comparisons among studies. Differential proteomic changes between conditions are often considered to be more reproducible than absolute abundances and may therefore provide a robust basis for cross-dataset integration. However, the systematic ability of differential change-based approaches to capture biologically meaningful relationships across heterogeneous datasets remains unclear.

**Results:** We developed a differential-change framework and applied it to public proteomic datasets. Pairwise contrasts were defined as differential proteomic profiles, and the concordance of up- and down-regulated proteins was quantified using odds ratios. Significant profile pairs were visualized as an integrative network. The treatment of anti-cancer drug doxorubicin vs control (MCF-7) comparison emerged as a central hub, with breast cancer proteome profiles clustering around it and associating with tumor stage (p = 0.03). Enrichment analysis revealed overrepresentation of lipid- and cholesterol-related pathways.

**Availability and implementation:** The source code for proteome network integration is available at https://github.com/manakanishizaki/proteome-network-integration.git.

## 1 Introduction

Proteomics aims to comprehensively characterize proteins with protein abundances, modifications, and interaction networks in biological systems and plays a key role in identifying cellular mechanisms, disease processes, and new biomarkers (Aebersold and Mann, 2016). Advances in mass spectrometry (MS) have enabled simultaneous quantification of thousands of proteins with high sensitivity and accuracy (Birhanu 2023). Concurrently, large volumes of proteomic data generated across laboratories and analytical platforms have been deposited in public repositories. Major resources include PRIDE (Perez-Riverol *et al*., 2025), MassIVE (Choi *et al*., 2020), and jPOST (Okuda *et al*., 2025), members of the ProteomeXchange Consortium (Deutsch *et al*., 2026), as well as the Clinical Proteomic Tumor Analysis Consortium (CPTAC) (Edwards *et al*., 2015), which provides extensive cancer proteomics datasets. Additionally, initiatives such as quantMS (Dai *et al*., 2024) provide standardized workflows for reanalyzing public datasets and disseminating results.

Two principal quantification strategies are commonly employed in MS-based proteomics: label-free quantification (LFQ) (Bantscheff *et al*., 2012) and labeling-based approaches (Wang *et al*., 2024). LFQ estimates relative protein abundance from peptide intensities or spectral counts (Bantscheff *et al*., 2012) without using chemical tagging, facilitating analysis of large cohorts. In contrast, labeling-based methods, such as tandem mass tag (TMT) (Thompson *et al*., 2003) and isobaric tag for relative and absolute quantitation (Chong *et al*., 2006; Rauniyar and Yates 2014) combine multiple samples within a single MS run, reducing run-to-run variability and enabling precise relative quantification.

Despite these advances, technical variability remains a major barrier to cross-study comparison. Differences in instrumentation, sample preparation, labeling strategies, and computational pipelines restrict direct comparability of proteomic measurements (Bantscheff *et al*., 2012). Absolute protein abundance estimates are particularly susceptible to batch effects, complicating data reuse and cross-platform integration. Such variability impedes large-scale meta-analyses and limits full use of rapidly expanding public proteomic resources.

Ratio-based quantitative strategies have also been shown to improve cross-platform reproducibility in multi-omics studies (Zheng *et al*., 2024). Differential proteomic changes between two conditions are considered more robust to technical variability and more reproducible across analytical platforms. In metabolomics, differential-based integration methods, such as iDMET (Matsuta *et al*., 2022), have captured cross-study relationships by comparing differential profiles, enabling extraction of biological signals independent of measurement platforms.

In the present study, we extended this differential-based integration framework to proteomics. Through systematic analyses of large-scale public datasets, we aimed to identify biological relationships conserved across studies and platforms. This framework enables robust detection of disease-associated protein changes and shared response patterns while minimizing platform-dependent effects, providing a scalable strategy for cross-study integrative proteomics.

## 2. Materials and methods

### 2.1 Data sources

We used publicly available, label-based quantitative proteomic datasets to systematically evaluate concordance in the direction of protein expression changes across conditions and to analyze the resulting network structure and its biological and clinical relevance.

We used two large-scale proteomics repositories: quantMS (Dai *et al*., 2024) and CPTAC (Edwards *et al*., 2015), both providing labeled quantitative MS datasets. From quantMS, we retrieved labeled-quantification projects, totaling 36 projects and 2,371 experimental conditions. From CPTAC, we obtained 49 TMT-based projects comprising 5,260 conditions. A complete list of all project identifiers is provided in Supplementary Table S1.

### 2.2 Preprocessing and quality control

Prior to analysis, protein identifiers from quantMS and CPTAC were standardized (Fig. 1A) and subjected to quality control. In quantMS, entries containing multiple UniProt identifiers were split to ensure a one-to-one correspondence between protein IDs and data rows, and entries labeled *DECOY* or *CONTAMINANT* were removed. For UniProt-formatted identifiers (sp|Accession|`ProteinName`), only the protein name was retained, and species suffixes (e.g., _ `HUMAN`) were removed to standardize notation.

**Figure 1.**
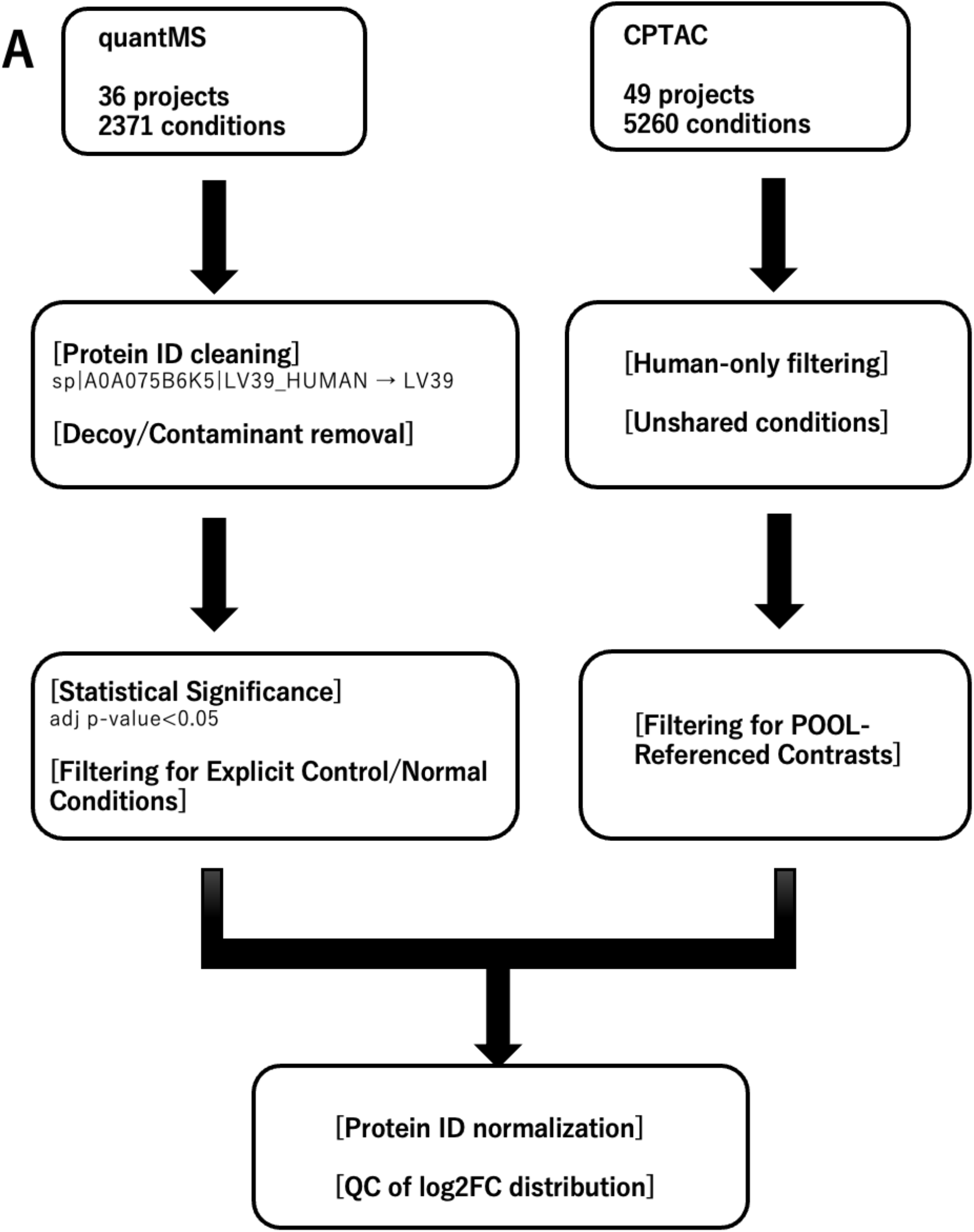
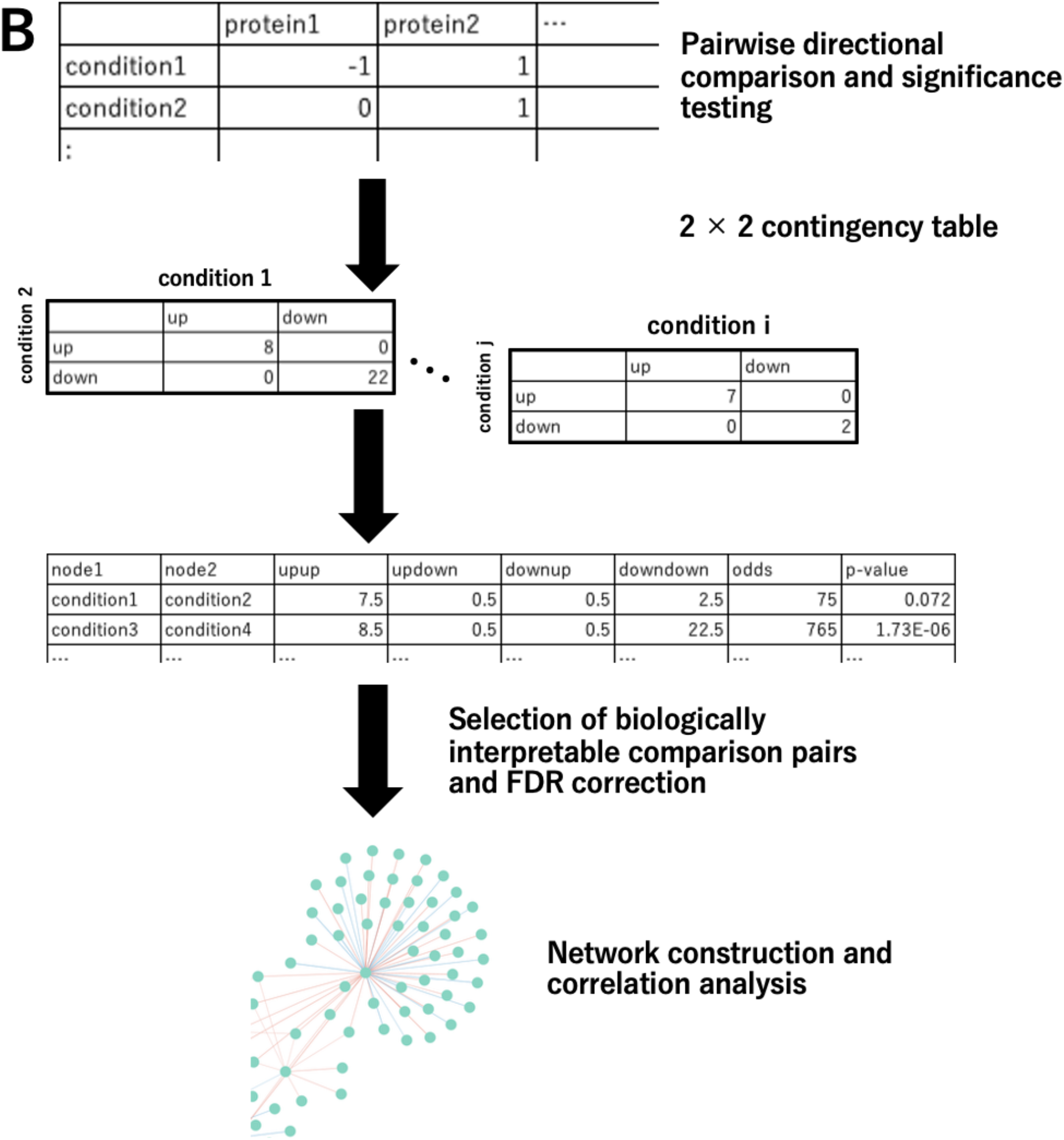
Workflow for identifier normalization and network construction. A) Overview of proteomic data collection and preprocessing. Proteomic datasets were obtained from quantMS (36 projects and 2,371 experimental conditions) and CPTAC (49 projects and 5,260 experimental conditions). In quantMS, protein identifiers were cleaned and standardized, including removal of decoy and contaminant entries. In CPTAC, analyses were limited to *Homo sapiens* proteins and conditions labeled “unshared.” After dataset-specific filtering, identifiers were normalized across resources, and the log2 fold-change (log2FC) distributions were evaluated prior to downstream analysis. B) Differential-based integration and network construction. For each condition, differential protein changes were encoded as directional signatures. Pairwise comparisons were performed to assess concordant and discordant directional changes, followed by statistical testing. Biologically interpretable pairs were selected using a false discovery rate (FDR) threshold and used to construct a network, enabling correlation analysis and identification of relationships across studies.

For CPTAC, only proteins annotated as *Homo sapiens* in the Organism field were included, and analyses were restricted to conditions labeled “unshared” to avoid interference from shared peptides and ensure more reliable quantification. To define significance thresholds, log_2_ fold-change (log2FC) distributions were examined across all conditions. Because log2FC values were largely within ±1, an absolute log_2_FC ≥ 1 was set as the cutoff. In quantMS, log2FC and multiple testing-adjusted p-values were available and used for filtering (|log2FC| ≥ 1 and adjusted p < 0.01). Conversely, because CPTAC data lacked p-values, only log2FC thresholds were applied.

### 2.3 Pairwise directional comparison and significance testing

Directional concordance of protein expression changes was evaluated for all condition pairs using directionality tables derived from quantMS and CPTAC. Proteins were classified as upregulated (log2FC ≥ 1) or downregulated (log2FC ≤ −1), and 2 × 2 contingency tables were constructed for each pair (Fig. 1b). Odds ratios and statistical significance were computed from upregulation and downregulation patterns. When necessary, odds ratios incorporated a Haldane–Anscombe continuity correction, and chi-square tests were used to assess independence of expression direction. Analyses were conducted for all condition pairs within each dataset, followed by extraction of biologically interpretable contrasts and application of a false discovery rate (FDR) correction.

### 2.4 Selection of biologically interpretable comparison pairs and FDR correction

Among all tested pairs, only biologically interpretable contrasts with consistent reference conditions were retained. In CPTAC, contrasts representing changes relative to pooled reference samples were selected, whereas in quantMS, contrasts with explicit references (e.g., control or normal) were included. Analyses were restricted to pairs from different projects; CPTAC projects reanalyzed in quantMS were excluded to preserve independence.

P-values obtained from directional concordance testing were adjusted using the Benjamini– Hochberg FDR procedure, and pairs with FDR <0.01 were deemed significant.

### 2.5 Network construction and correlation analysis

Networks were generated from filtered contrasts in quantMS and CPTAC. Nodes represent comparison conditions, and edges connect pairs showing significant directional concordance in protein expression (FDR < 0.01). Network construction and visualization were performed using Cytoscape (Shannon. *et al*., 2003). From the resulting network, the pair with the highest odds ratio was selected to determine whether directional concordance reflected consistent expression changes. Log2FC values for proteins quantified in both conditions were extracted, Pearson correlation coefficients were calculated, and scatter plots were generated to evaluate agreement between conditions and consistency with network-derived association strength.

### 2.6 Hierarchical clustering of expression profiles across conditions

To assess condition similarity, log2FC values for proteins commonly quantified across contrasts were integrated into a matrix, focusing on quantMS-derived contrasts and their connected CPTAC counterparts. Infinite values were treated as missing, and missing log2FC values were imputed as zero, assuming no detectable change in the absence of quantification. This approach was adopted to avoid introducing artificial correlations and to maintain interpretability across contrasts, given that the analysis focuses on relative expression changes rather than absolute abundances. Similarity among conditions and proteins was evaluated using hierarchical clustering with Euclidean distance and Ward’s linkage, applied to rows (proteins) and columns (contrasts) to visualize expression patterns.

### 2.7 Association with clinical variables and functional enrichment analysis

Associations between contrast clusters and clinical categorical variables were evaluated using chi-square tests. Analyses were conducted using CSV files containing patient clinical annotations, with the following variables considered: ethnicity, race, primary diagnosis, tissue or organ of origin, tumor grade, tumor stage, disease type, and primary site. Samples labeled “Not Reported” or “Unknown” were excluded.

To support biological interpretation of clustered proteins, functional enrichment analysis was performed via Metascape (Zhou *et al*., 2019) based on Gene Ontology (GO) Biological Process terms (Ashburner *et al*., 2000) and Kyoto Encyclopedia of Genes and Genomes (KEGG) pathway (Kanehisa and Goto, 2000) annotations. Subtype-specific analyses were not performed because certain cancer subtype annotations reported in the original CPTAC publications were unavailable in the clinical datasets analyzed.

## 3. Results

### 3.1 Establishment of the study workflow

We developed a standardized analytical workflow to integrate heterogeneous proteomic datasets and construct a differential change-based condition network (Fig. 1). Proteomic data were collected from quantMS (Dai *et al*., 2024) and CPTAC (Edwards *et al*., 2015) and subjected to dataset-specific preprocessing and filtering. Protein identifiers were cleaned and standardized, including removal of decoy and contaminant entries and restriction to human proteins where appropriate. After filtering, protein identifiers were normalized across resources, and log2FC distributions were assessed to confirm comparability before downstream analyses.

Unlike conventional approaches that primarily focus on within-dataset analyses, our framework enables cross-study integration by capturing condition-level relationships based on directional concordance of differential protein changes.

For each experimental condition, differential protein changes were encoded as directional signatures. Pairwise comparisons between conditions were performed to quantify concordant and discordant directional changes, followed by statistical testing and FDR-based selection of biologically interpretable condition pairs. These retained pairs were used to construct a condition-level network, enabling analysis of relationships among experimental conditions across independent studies.

### 3.2 Quality control and significant proteins

The analyzed projects encompassed diverse contrasts, including drug-treated versus control samples and tumor versus normal tissues. The quantMS and CPTAC datasets identified 14,508 and 15,328 proteins, respectively, with 6,303 proteins detected in both resources (Fig. 2A).

**Figure 2.**
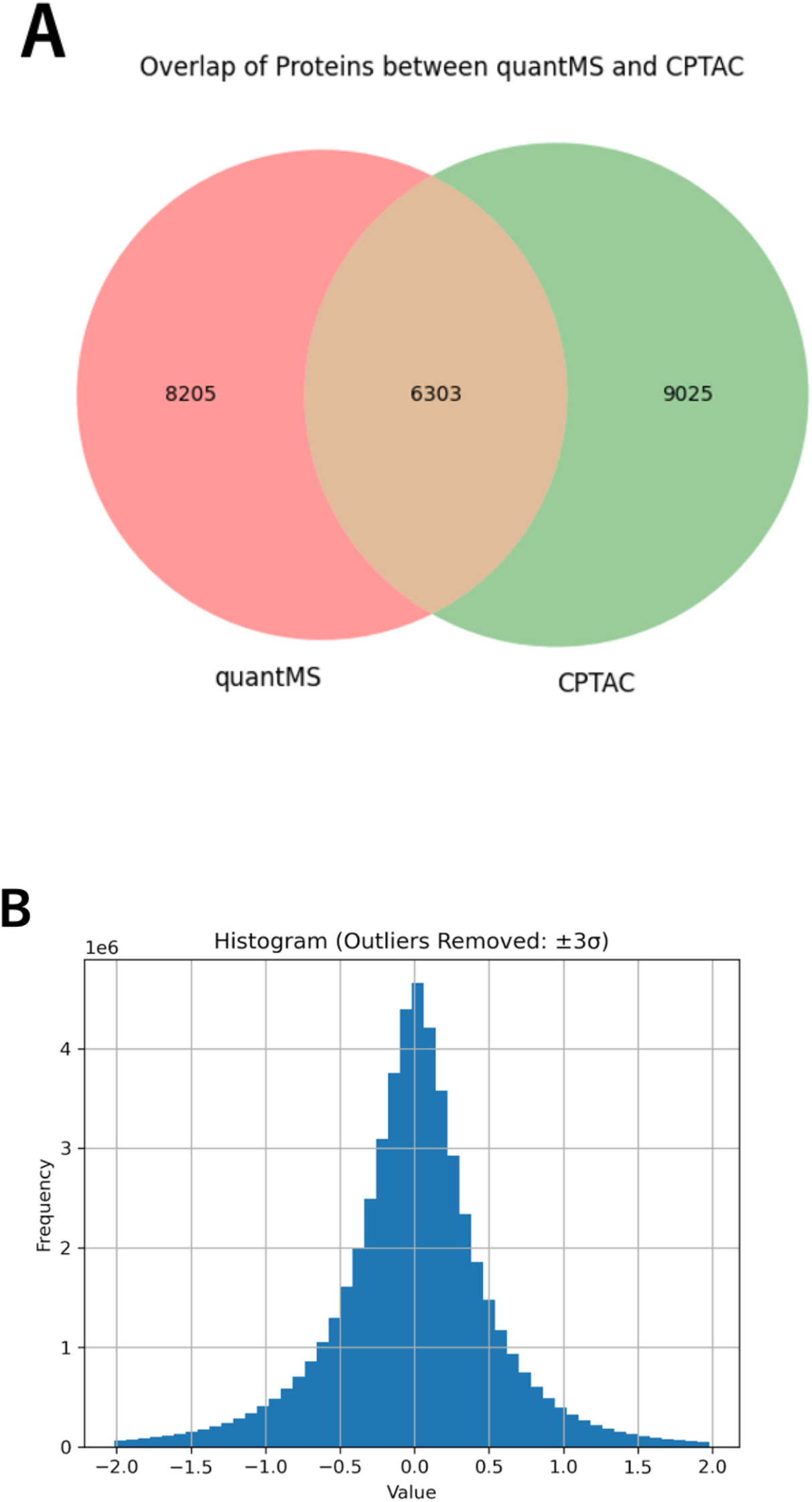
Dataset statistics. A) Overlap of proteins identified in quantMS and CPTAC. Venn diagram shows overlap between the 14,508 and 15,328 proteins detected in quantMS and CPTAC, respectively, with 6,303 shared proteins identified. B) Distribution of log2FC values in quantMS and CPTAC. Histograms show log2FC distributions across all datasets. Values exceeding ±3 standard deviations from the mean were excluded from visualization to improve clarity but retained in downstream analyses and quantitative datasets.

To characterize differential expression, percentile-based metrics were used to summarize global fold-change distributions. Across projects, most log2FC values fell within ±1 (5 th–95 th percentile: −1.03 to 0.92; Fig. 2B and Supplementary Fig. S1), indicating that most proteins exhibited limited changes while a minority showed large expression shifts. Based on this distribution, proteins with at least a twofold change (|log2FC| ≥ 1) were defined as significantly altered. These stable global fold-change patterns support subsequent pairwise comparisons and network analyses.

### 3.3 Initial network construction and observations

To compare fold-change directionality across quantMS and CPTAC, only contrasts with consistent, interpretable reference conditions were retained. In CPTAC, contrast annotations were heterogeneous: many contrasts not explicitly formatted as “A/B” were inferred to represent changes relative to pooled references (POOL), whereas explicitly annotated “A/B” comparisons often involved non-POOL references, reference–reference comparisons (e.g., QC–QC, bridge–bridge, or CR–CR), or sample–sample contrasts lacking a shared reference. Because these formats were unsuitable for directional analysis, they were excluded using predefined criteria.

After filtering, 463 CPTAC contrasts were removed, leaving 4,797 POOL-referenced contrasts. In quantMS, contrasts lacking explicit reference conditions were excluded, yielding 28 contrasts for downstream analysis. The resulting initial condition-level network comprised 639 nodes and 1,040 edges (Supplementary Fig. S2).

The highest odds ratio was observed between “PDC000126_normal–ovarian serous carcinoma” (quantMS-derived) and “PDC000125_CPT0026540001 Unshared Log Ratio” (CPTAC-derived). PDC000126 represents a reanalysis of the CPTAC UCEC Discovery Study phosphoproteome, whereas PDC000125 corresponds to a proteome dataset from the same CPTAC study (Dou *et al*., 2023). Although they were assigned different project identifiers, these datasets originate from the same underlying CPTAC study and differ primarily in the measured molecular layer.

Despite reprocessing through quantMS, PDC000126 showed the strongest directional concordance with the original CPTAC proteome dataset. This finding indicates that differential-change patterns remain reproducible across analytical pipelines when derived from the same biological source, supporting the validity of our approach. Because this strongest association involved datasets from the same CPTAC project, CPTAC studies reanalyzed within quantMS were excluded from subsequent analyses to focus on biologically informative relationships among truly independent datasets across databases.

### 3.4 Network established after removal of reanalyzed CPTAC projects

After excluding CPTAC projects that had been reanalyzed on quantMS, the refined network contained 287 nodes and 357 edges (Fig. 3). Each edge represented a condition pair with significant directional concordance based on the chi-square test. The network topology revealed a subset of highly connected nodes. These hub nodes corresponded to experimental conditions exhibiting directional concordance with multiple other conditions and were frequently associated with relatively large differential changes. Major hub nodes and their degrees are summarized in Table 1.

**Table 1.**
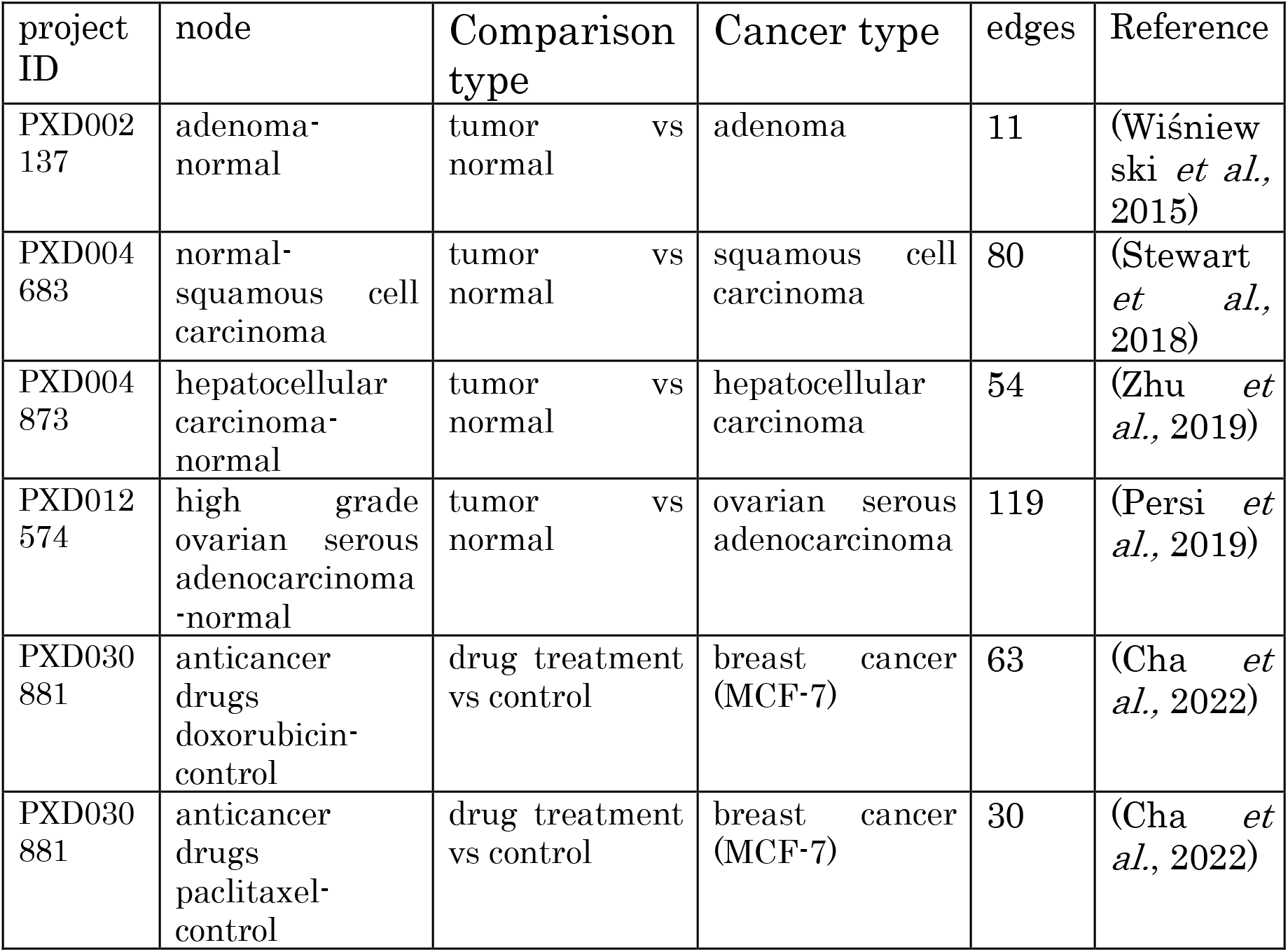
Hub nodes identified in the integrated proteomic network derived from quantMS and CPTAC data. Each node represents a comparison between control and treated conditions within the same biological context, e.g., tumor vs. normal tissue or drug treatment vs. control. Number of edges indicates node degree, reflecting the number of significant correlations with other nodes in the network. Associated cancer types and project identifiers are provided.

**Figure 3.**
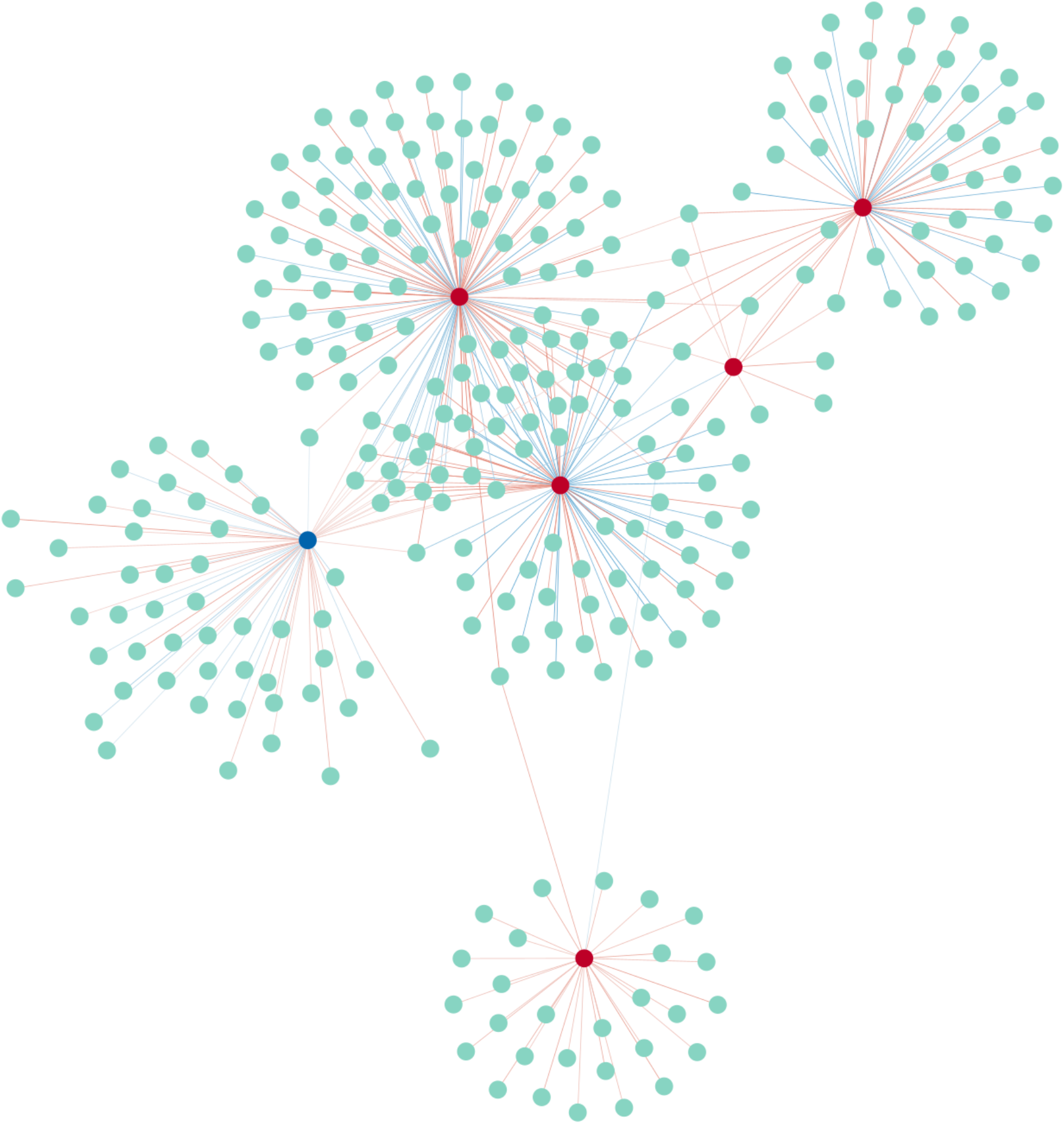
Integrated network constructed after excluding CPTAC projects reanalyzed by quantMS. The network was generated after removing CPTAC-derived projects reanalyzed by quantMS and consists of 287 nodes and 357 edges. Edges represent condition pairs with statistically significant associations based on chi-square tests with FDR correction (q < 0.01). Edge colors indicate the direction of association (red for positive and blue for negative). Hub nodes are highlighted in red, and the node of particular interest (PXD030881_anticancer doxorubicin–control) is highlighted in blue.

### 3.5 Correlation analysis of the highest odds-ratio pair

For the pair with the highest odds ratio, correlation analysis was performed between ovarian cancer (PXD012574_high grade ovarian serous adenocarcinoma–normal) and uterine cancer (PDC000439_CPT0127980003 Unshared Log Ratio) using the log2FC values of commonly quantified proteins (Fig. 4).

**Figure 4.**
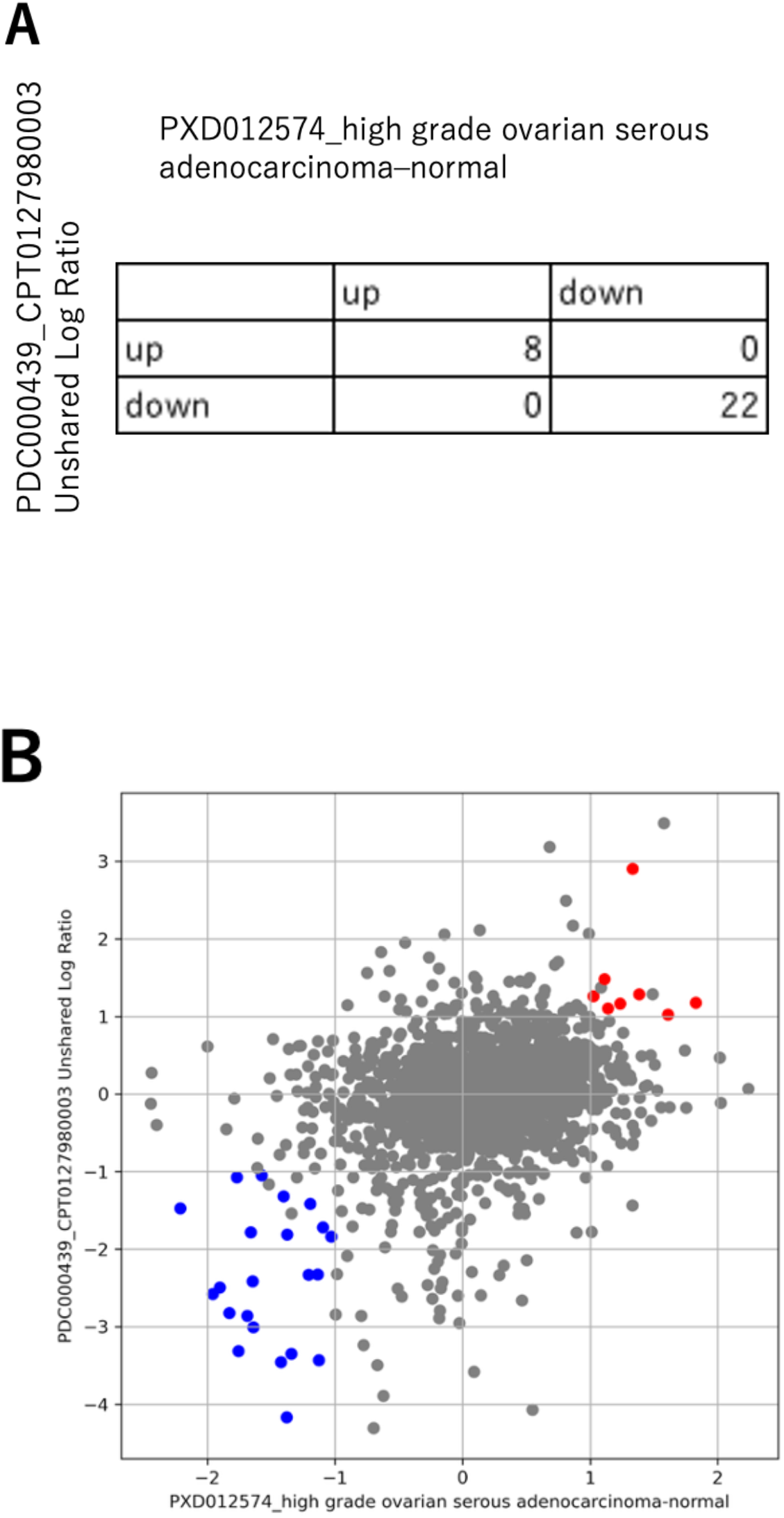
Comparison of log2FC values for protein expression between ovarian and uterine cancers. A) Contingency table (2 × 2) summarizing proteins with concordant increases or decreases across conditions. Association was assessed using the chi-square test, yielding a log2 odds ratio of 7.65 and a p-value of 1.7 × 10^−6^. B) Protein log2FC compared between ovarian cancer (PXD012574_high grade ovarian serous adenocarcinoma–normal) and uterine cancer (PDC000439_CPT0127980003 Unshared Log Ratio). Proteins exceeding the predefined threshold (|log2FC| ≥ 1) are highlighted: red, upregulated; blue, downregulated.

Cross-tabulation revealed that 8 and 22 proteins were significantly upregulated and downregulated, respectively, under both conditions, whereas no proteins showed significant changes in opposite directions. Across all quantified proteins, the Pearson correlation coefficient was 0.318, indicating moderate overall association. In contrast, restricting analysis to significantly altered proteins increased the coefficient to 0.801, demonstrating strong concordance in expression direction.

Although representing distinct tumor types, ovarian and uterine cancers exhibited similar proteomic alteration patterns. This similarity is biologically plausible because both tumors are epithelial in origin and share oncogenic processes, including PI3K/AKT and MAPK signaling, hormone responsiveness, metabolic reprogramming, and cytoskeletal remodeling (Cheaib 2015; Aseervatham 2020). These shared molecular features likely underlie the strong directional concordance observed among significantly altered proteins.

### 3.6 Biological interpretations

We systematically evaluated relationships among experimental conditions from multiple proteomics databases using network analysis and hierarchical clustering of log2FC profiles. As a representative case, we examined a hub node corresponding to the breast cancer cell line MCF-7 treated with the anticancer drug doxorubicin (PXD030881_anticancer doxorubicin–control). This condition was selected because it exhibited numerous significant similarities to other conditions and represented a well-defined drug-versus-control contrast.

Hierarchical clustering centered on this doxorubicin–control condition identified one clearly defined major cluster of breast cancer–related experimental conditions, whereas other breast cancer contrasts were more dispersed (Fig. 5). Proteins within the major cluster showed coherent expression patterns, indicating shared molecular features across conditions.

**Figure 5.**
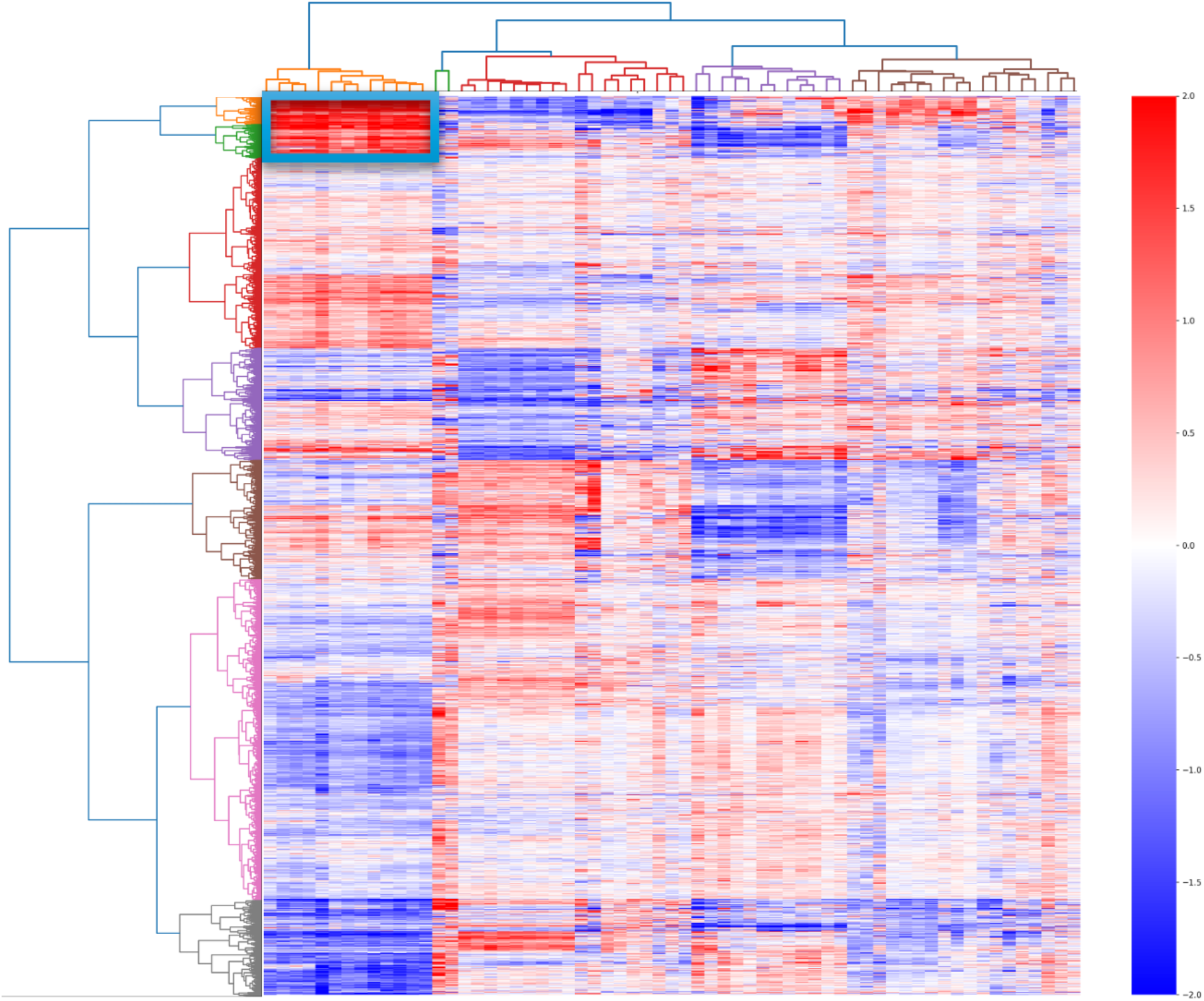
Heatmap of CPTAC-derived experiments associated with the quantMS-derived doxorubicin–control hub. Heatmap summarizes the protein expression patterns of CPTAC-derived experimental conditions linked to the quantMS-derived doxorubicin–control comparison. Proteins (rows) and conditions (columns) were hierarchically clustered, with dendrograms shown on the left and top, respectively. Leftmost condition cluster corresponds to the subset of breast cancer samples that was analyzed further.

Integration with clinical metadata indicated that samples in the major cluster tended to have more advanced tumor stages (chi-square test, p = 0.0386). Although the effect size was modest, this finding suggests a potential link between proteomic patterns and tumor progression.

Functional enrichment analysis of proteins upregulated in the major cluster revealed significant enrichment of pathways related to lipid metabolism, extracellular matrix (ECM) remodeling, and tissue repair (Supplementary Fig. S3). These processes have previously been implicated in breast cancer progression and tumor microenvironment remodeling (Zhao *et al*., 2017; Tang *et al*., 2022; Krug *et al*., 2020), supporting the biological relevance of the observed cluster structure.

Collectively, these findings demonstrate that the directional consistency–based network and clustering framework employed in this study can extract biologically and clinically interpretable patterns from heterogeneous proteomics datasets across independent resources.

## 4 Discussion

We developed and applied a directionality-based differential-change framework to systematically assess relationships among experimental conditions across heterogeneous proteomic datasets. Hierarchical clustering centered on the doxorubicin–control profile revealed one clearly defined major cluster and several heterogeneous groups, demonstrating the framework’s capacity to capture coherent condition-level groupings across independent studies.

Conditions within the major cluster were relatively enriched for advanced tumor stages, with a modest but statistically significant association. Upregulated proteins were enriched for pathways related to lipid metabolism and ECM remodeling, consistent with prior reports in breast cancer biology (Zhao *et al*., 2017; Tang *et al*., 2022), indicating that the network structure reflects biologically interpretable patterns present in large-scale heterogeneous proteomic data. In contrast, downregulated proteins showed modest overrepresentation of core cellular processes, including translation, endoplasmic reticulum protein processing, and DNA replication. This asymmetry suggests that the framework preferentially captures condition-specific regulatory programs rather than broadly conserved housekeeping functions.

For cross-dataset integration, protein-level changes were defined based on fold-change directionality using a |log2FC| ≥ 1 threshold. This strategy reduces sensitivity to platform-dependent differences in quantitative scale and normalization. Consequently, fold-change magnitude was not explicitly incorporated into network construction or similarity assessment. While this approach mitigates variability arising from differences in experimental platforms and data-processing pipelines, it may reduce sensitivity to subtle but consistent changes or biological effects primarily reflected in fold-change magnitude.

Overall, the proposed framework recovers biologically and clinically coherent patterns across independently generated proteomic datasets. Consistent differential-change patterns were observed despite differences in measurement platforms and analytical pipelines, supporting the robustness of this approach for exploratory cross-dataset integration. The identified relationships should therefore be regarded as hypothesis-generating findings that provide a foundation for future experimental validation and targeted investigations.

Several limitations of the present study should be noted. First, integration across proteomic datasets relied on protein identifiers as reported in the original studies. Although efforts were made to harmonize identifiers, inconsistencies in protein naming conventions, accession numbers, and annotation versions across datasets may have resulted in incomplete matching. Such discrepancies could lead to an underestimation of shared differential signals between studies and potentially obscure biologically relevant relationships. Further standardization of protein identifiers and systematic mapping across database versions would likely improve the accuracy and sensitivity of cross-dataset network analyses.

Second, our analysis was restricted to publicly available labeled quantitative proteomic datasets. While labeled approaches provide relatively consistent quantification within studies, label-free quantitative data is more abundant in the repositories. As proteomic repositories continue to expand and data-sharing practices improve, incorporating a broader range of studies will increase statistical power and enhance the discovery of novel condition-level relationships.

Future work will extend the present framework to label-free quantitative proteomic datasets. A substantial number of label-free datasets are publicly available, and adapting the directionality-based integration strategy to these data will allow evaluation of its robustness across different quantification platforms. Such expansion will enable larger and more diverse network constructions, potentially uncover additional biologically meaningful cross-study relationships and further establish the general applicability of the proposed framework.

## 5 Conclusion

In this study, we applied a differential-change–based integrative framework to proteomic data and systematically evaluated shared proteomic alteration patterns across diverse experimental conditions from quantMS and CPTAC. By emphasizing expression directionality, we demonstrated that biologically coherent relationships can be consistently extracted from large-scale datasets generated using different platforms and analytical pipelines. Analysis centered on the doxorubicin–control profile identified a distinct cluster of breast cancer samples associated with tumor stage and lipid metabolism–related pathways. These findings align with prior breast cancer studies and support the validity of the proposed approach for capturing biologically meaningful patterns in heterogeneous proteomic data. Overall, this research provides a practical, robust framework for cross-dataset proteomic data integration without relying on absolute expression levels or individual-protein statistical significance. The results provide a foundation for exploratory analyses and hypothesis generation, facilitating future validation and integrative proteomics research. We expect that the proposed strategy will encourage broader reuse of publicly available proteomic datasets and foster new insights through systematic cross-study integration.

## Author contributions

Manaka Nishizaki (Data curation [lead], Formal analysis [lead], Investigation [lead], Methodology [lead], Software [lead], Validation [lead], Visualization [lead], Writing—original draft [lead]), Norie Araki (Formal analysis [supporting], Methodology [supporting], Supervision [supporting], Writing—review & editing [equal]), and Shin Kawano (Conceptualization [lead], Funding acquisition [lead], Methodology [supporting], Project administration [lead], Supervision [lead], Writing—original draft [supporting], Writing— review & editing [equal]).

## Conflict of interests

None declared.

## Funding

This work was supported by Database Integration Coordination Program, Japan Science and Technology Agency (grant number: 23810525 and 23810055).

## Data availability

The datasets analyzed in this study are publicly available. Proteomics data were obtained from the quantMS repository and the CPTAC data portal. The identifiers and accession numbers for all datasets used in this study are provided in Supplementary Table S1. The codes used by this research is available at https://github.com/manakanishizaki/proteome-network-integration.git. All data and codes are accessible without restriction.

## Notes

### Competing Interest Statement

The authors have declared no competing interest.

