## Supplementary data for "Network-based integration of cross-dataset proteomic profiles using fold-change directionality"

Supplementary Table S1. Proteomic projects included in this study.

CPTAC TMT10, CPTAC TMT11, and quantMS projects analyzed in this study are summarized. In quantMS, some datasets sharing the same ProteomeXchange accession (e.g., PXD023508) are listed separately when associated with substantially different experimental condition labels (e.g., disease, mutation carrier, and phenotype). Although derived from the same study, these datasets are treated as distinct entries.

| CPTAC TMT10 | CPTAC TMT11 | quantMS |
|---|---|---|
| PDC000110 | PDC000180 | PDC000111 |
| PDC000116 | PDC000198 | PDC000126 |
| PDC000118 | PDC000203 | PXD000672 |
| PDC000120 | PDC000204 | PXD002137 |
| PDC000124 | PDC000221 | PXD002395 |
| PDC000125 | PDC000234 | PXD003497 |
| PDC000127 | PDC000270 | PXD003539 |
| PDC000129 | PDC000315 | PXD004683 |
| PDC000153 | PDC000316 | PXD004684 |
| PDC000154 | PDC000317 | PXD004691 |
| PDC000219 | PDC000325 | PXD004873 |
| PDC000242 | PDC000358 | PXD010429 |
| PDC000248 | PDC000360 | PXD012574 |
| PDC000295 | PDC000362 | PXD014145 |
| PDC000296 | PDC000402 | PXD014414 |
| PDC000309 | PDC000408 | PXD014943 |
| PDC000310 | PDC000410 | PXD018830 |
| PDC000311 | PDC000434 | PXD020109 |
| PDC000327 | PDC000439 | PXD020248 |
| PDC000329 | PDC000440 | PXD021394 |
| PDC000356 | PDC000446 | PXD022992 |
| PDC000393 | PDC000447 | PXD023423 |
| PDC000398 | PDC000464 | PXD023508-disease |
| PDC000400 | PDC000477 | PXD023508-mutation-carrier |
| | PDC000514 | PXD023508-phenotype |
| | | PXD025560 |
| | | PXD025864 |
| | | PXD027008 |
| | | PXD027817 |
| | | PXD028251 |
| | | PXD028618 |
| | | PXD030671 |
| | | PXD030881 |
| | | PXD032263 |
| | | PXD033169 |

Supplementary Figure S1. Distribution of log2FC values by proteomic project.

Box plots show log2FC distributions for individual quantMS and CPTAC projects. Values exceeding ±3 standard deviations from the project-specific mean were excluded from visualization for clarity but retained in downstream analyses and quantitative datasets.

Supplementary Figure S2. Integrated network prior to excluding CPTAC datasets reanalyzed in quantMS.

Network contains 639 nodes and 1,040 edges. Edge colors indicate correlation direction: red, positive; blue, negative. Network was generated before removing CPTAC-derived data included in quantMS. Hub nodes are highlighted in red, whereas nodes of interest (from PDC000126) are shown in blue.

Supplementary Figure S3. Functional enrichment analysis of representative protein clusters.

GO and KEGG pathway enrichment analyses were performed using Metascape for representative protein clusters showing increased (A) or decreased (B) abundance. Enriched terms are ranked by according to statistical significance (–log10 P value).

### Supplementary Figure S1.

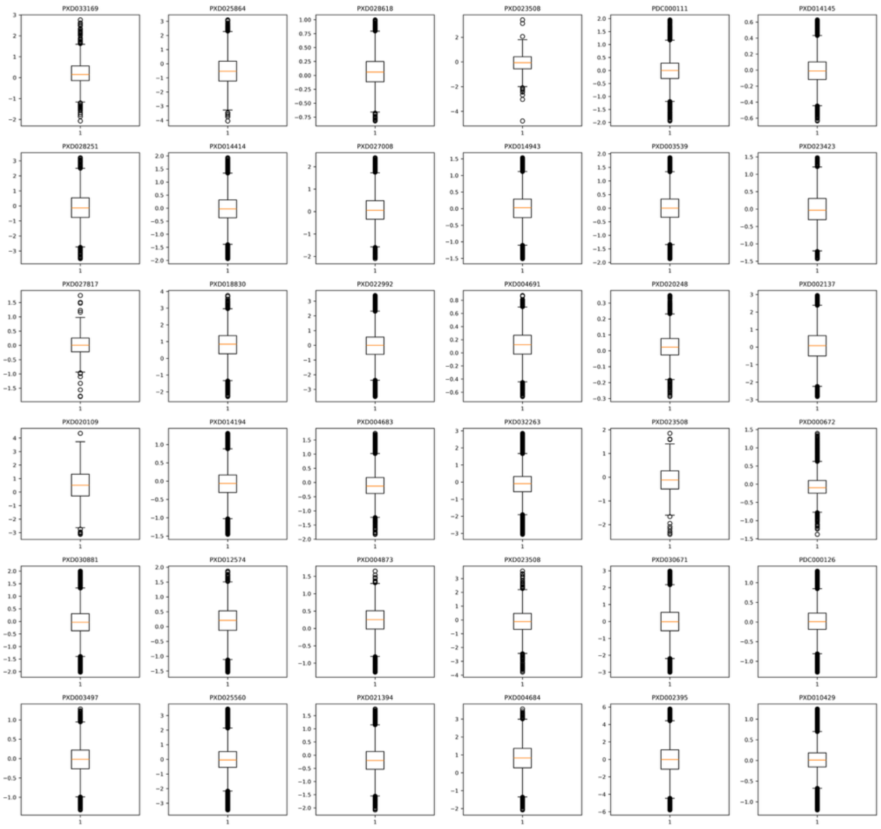

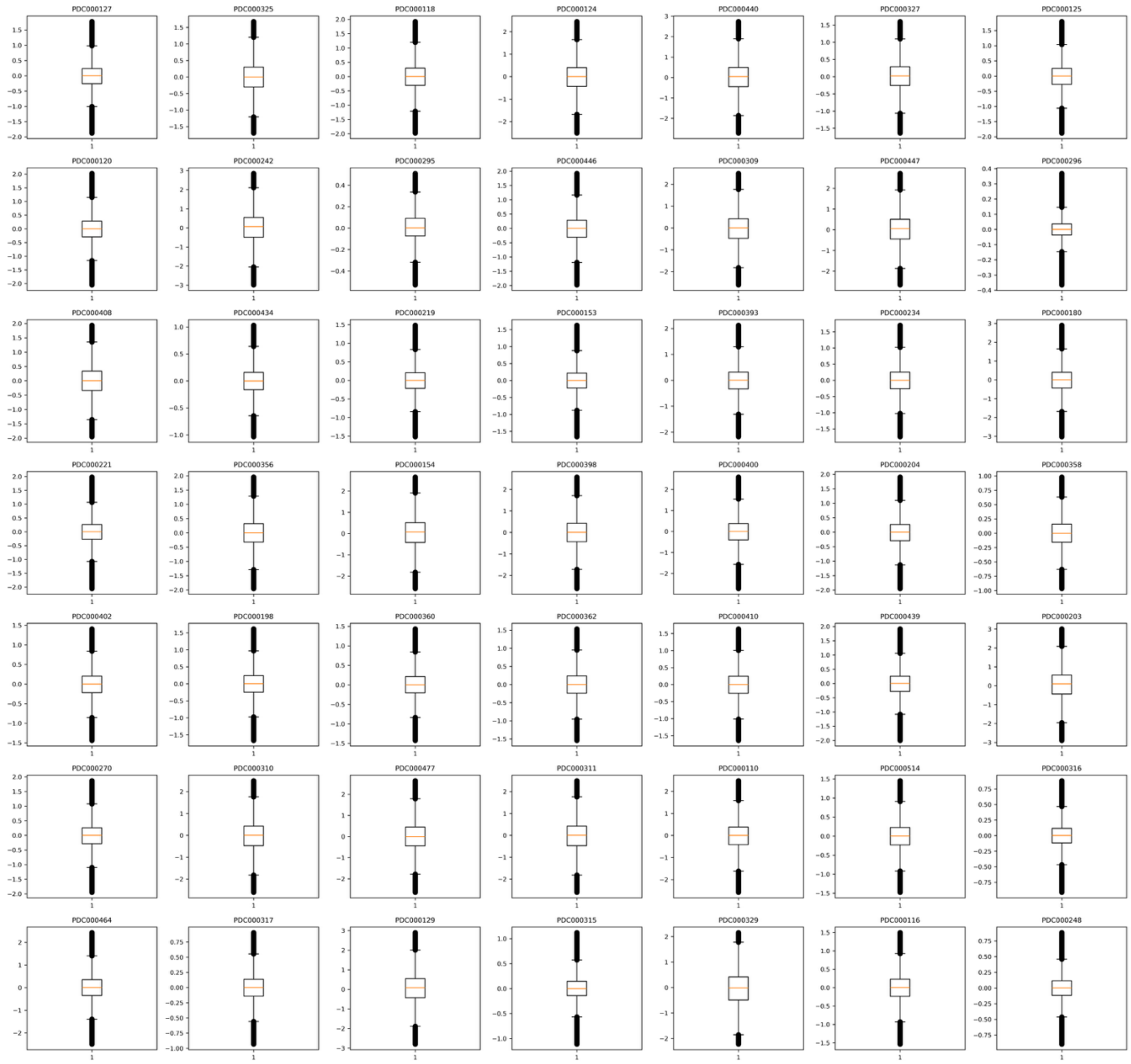
PDC000127
PDC000325
PDC000118
PDC000124
PDC000440
PDC000327
PDC000125
PDC000120
PDC000242
PDC000295
PDC000446
PDC000309
PDC000447
PDC000296
PDC000408
PDC000434
PDC000219
PDC000153
PDC000393
PDC000234
PDC000180
PDC000221
PDC000356
PDC000154
PDC000398
PDC000400
PDC000204
PDC000358
PDC000402
PDC000198
PDC000360
PDC000362
PDC000410
PDC000439
PDC000203
PDC000270
PDC000310
PDC000477
PDC000311
PDC000110
PDC000514
PDC000316
PDC000464
PDC000317
PDC000129
PDC000315
PDC000329
PDC000116
PDC000248

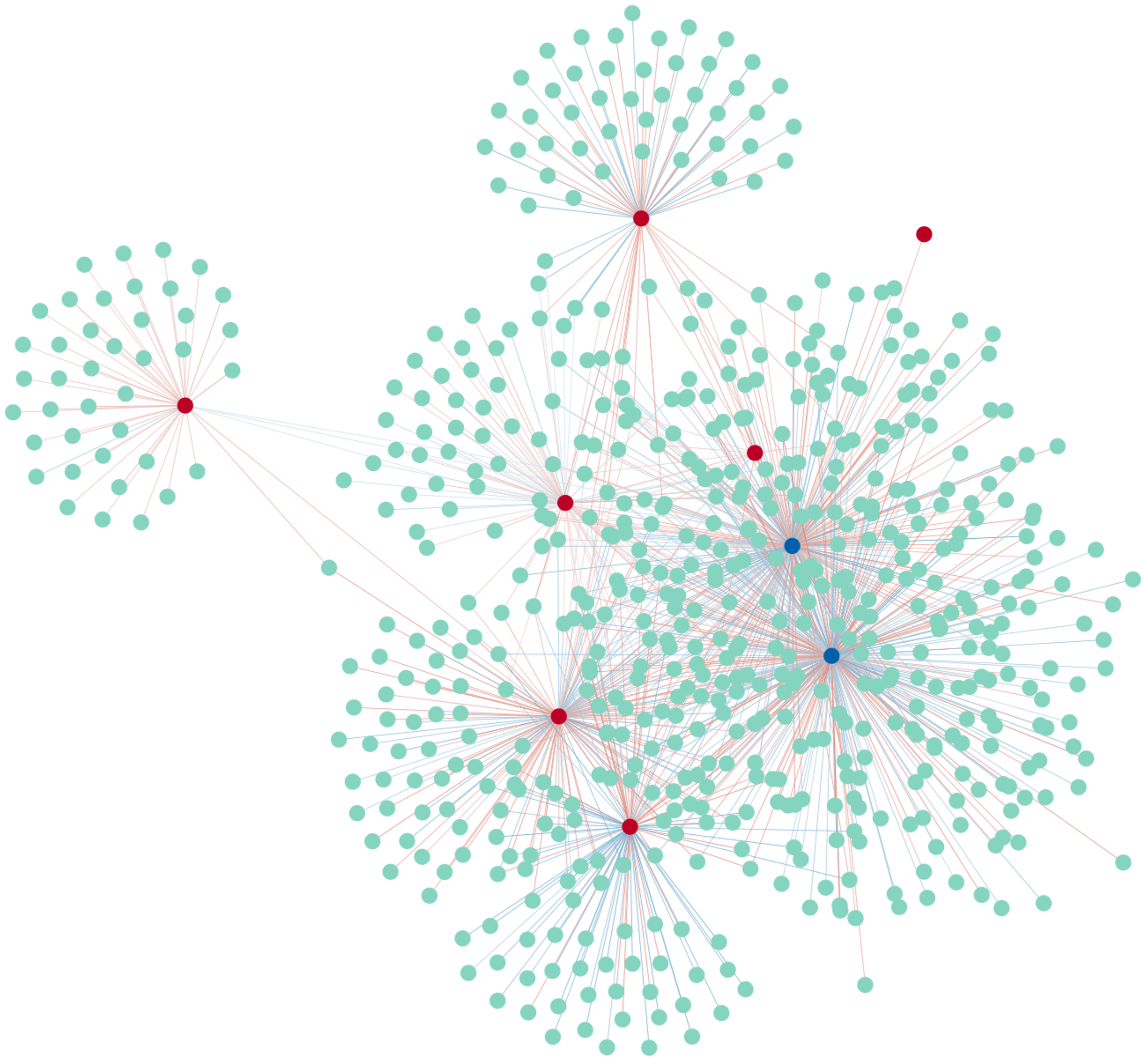

**Supplementary Figure S2.**

A

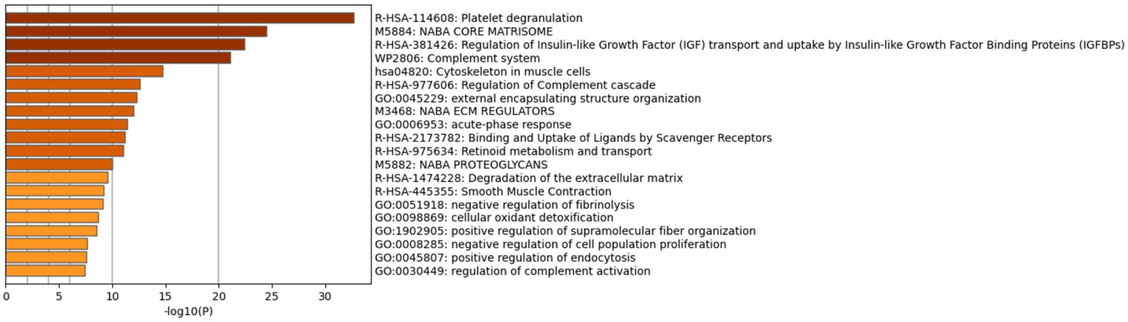


B

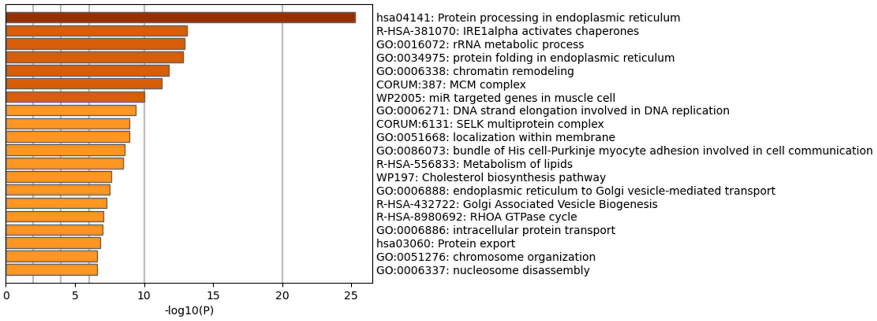


**Supplementary Figure S3**
